# Characterization of network hierarchy reflects cell-state specificity in genome organization

**DOI:** 10.1101/2022.08.03.502724

**Authors:** Jingyao Wang, Yue Xue, Yueying He, Hui Quan, Jun Zhang, Yi Qin Gao

## Abstract

Dynamic chromatin structure acts as the regulator of transcription program in crucial processes including cancer and cell development, but a unified framework for characterizing chromatin structural evolution remains to be established. Here, we performed graph inferences on Hi-C datasets and derived the chromatin contact networks (CCNs). We discovered significant decreases in information transmission efficiencies in CCNs of colorectal cancer (CRC) and T-lineage acute lymphoblastic leukemia (T-ALL) compared to corresponding normal controls through graph statistics. Through network embedding in Poincaré disk, the hierarchy depths of CCNs from CRC and T-ALL patients were found to be significantly shallower compared to their normal controls. A reverse trend of CCN change was observed for the transition from embryo to differentiated primary tissues. During early embryo developments of both human and mouse, the hierarchy of embedded chromatin structure gradually establishes. Additionally, we found tissue-specific conservation of hierarchy order in normal CCNs, which was disturbed in tumor. Our findings uncover the cell-state related hierarchical characteristics in chromatin structure, which reveal chromatin loci that play important roles in stabilizing the cellular conditions.

## INTRODUCTION

In three-dimensional space, chromatin structure is organized in a nested hierarchy, in a way that higher-grade entities are composed of lower-grade entities in a nested way^1^. At the nucleus scale, chromatin consists of relatively independent chromosome territories^2^. At the micron scale, chromatin is divided into heterochromatin and euchromatin with different sequential characteristics, openness, epigenetic landscapes and transcription activities^3,4^. Heterochromatin and euchromatin are also manifested by compartments in Hi-C matrix^5^. At the nanometer scale, dynamic clusters with a range of physical sizes are formed, corresponding to nested TADs in Hi-C matrix^6-8^. To decipher hierarchically organized TADs, graph theory was found to be useful^9,10^.

While chromatin structure has a nested hierarchy, to the best of our knowledge, its flow hierarchy has not been investigated. Flow hierarchy describes an architecture where information flows from higher-grade nodes to lower-grade ones^11^. Chromatin contact networks have a scale-free pattern, which is characterized by the power-law distribution of node degrees^12^. Power-law phenomena in nature imply existence of self-organization patterns and hidden hierarchy^13^. In scale-free networks, a few nodes have strong influences over others, thus have high flow-hierarchy statuses^14^. Networks with flow hierarchy are characterized by strong clustering, high heterogeneity and frequent occurrence of bridge edges which link network communities and guarantee efficient routing of signals^15^. Real-life complex networks perform information transmission tasks and are abstracted of graphs describing relationships (edges) between entities (nodes). Graph theory has been widely used in deciphering biological networks including protein interaction and gene expression networks^16^, where dynamic information transmission is vital for maintaining functionalities^17,18^. From the network view, chromatin architecture is also integrated with function of genetic information transmission. Instead of maintaining a static configuration, chromatin structure is significantly dynamic^19^. A large and diverse set of gene regulatory elements in chromatin are found to regulate transcription through long-range interactions. For example, promoter-enhancer interactions regulate dynamic activation of genes^20^, while insulators and repressors play the opposite roles^21^. With these annotated functional contacts acting as bridging edges, chromatin functions as a complex dynamic network transmitting signals vital for cell viability. The correspondence of chromatin-fragment interaction and the protein-protein interaction demonstrated that 3D genome contributes to protein interactome which directly exercises biological functions^22,23^. In addition, noncoding loci without any annotations are indispensable for stability of chromatin structure and maintaining cell identities, i.e., targeted deletion of chromatin loci close to many loci in space can computationally impose the network structural property of a cancer cell onto a normal cell^12^.

Chromatin structure evolves during crucial life processes including cancer and cell development. Although extensive studies have attributed oncogenesis to gene mutations^24-27^, targeting chromatin structure for diagnosis and therapy has increasingly gained attention^28,29^. Conventional microscopic observation of aberrant chromatin was used for cancer diagnosis^30^. Gradual changes brought by heterochromatin de-compaction through developmental stages of multiple cancers were observed using microscopy^31^. The recently developed Hi-C technology allows detailed inspection of aberrant chromatin structure in pathological tissues. Radial re-organization of compartment A/B in nucleus was observed in CRC^32^ and attenuation of long-range chromatin interactions in Hi-C was seen in multiple cancer cell-lines compared to normal cells^33^. Pathological cancer tissues were observed to be accompanied by disturbances in compartments and TAD structures^32,34,35^.

On the other side, similarity between embryonic stem cells and cancer cells has also attracted attention^36-38^. The shared characteristics by embryonic stem cells and cancer cells include unlimited proliferation and high cellular plasticity^39,40^. Critical pathways including Wnt and Notch signaling pathways are associated with both embryonic development and cancer progression^41,42^. Certain gene expression patterns in early embryonic development is re-triggered in cancer^43^. However, the similarity in chromatin structure between embryo tissues and pathological cancer tissues has not been extensively studied. During early embryo development, chromatin architecture is re-organized from a relaxed state to a mature one, reflected by establishment of compartments and TADs^44^. In lineage specifications, establishment of long-range promoter-enhancer contacts are correlated with corresponding gene activation^45^. Significant structural changes in chromatin in cancer and cell development underline the importance of genome architecture for maintaining stable cell identities^33,46-48^.

With the rapid development of experimental techniques for detecting chromatin structure, computational methods for deciphering chromatin structure are also emerging. However, existing methods for modeling chromatin structure are mainly limited in three-dimensional Euclidean space, which overlooks its functional behavior as a hierarchical network^12,20,21^. Through feature extraction, symbolic data such as texts and networks can be transformed into sets of vectors with representation learning, such a process is called embedding^49-51^. Embeddings of complex networks and multi-relational data have achieved great success in completing missing data and extracting information^52-54^. To embed a network is to map the network nodes to vectors in a high-dimensional space with maximally preserved network connectivity manifested by distances. Equipped with a negative curvature, hyperbolic space can pack in more surface area within a certain radius than its Euclidean counterparts with a zero-curvature. Since the geometry behind hierarchical networks is hyperbolic rather than Euclidean^15^, to embed a hierarchical network into hyperbolic space can preferably reserve its structural information, whereas embedding in Euclidean space may lead to distortion due to incompetent representation ability^55,56^.

Among several isoforms of hyperbolic space, Poincaré disk is superior in its differentiable distance formula with easy implementation of gradient descent algorithms as well as in its simple constraints^55^. Poincaré disk is a projected hyperbolic space on a unit circle where perimeter and area grow exponentially with the radius. Poincaré embedding allows natural characterization of hierarchical complex networks from massive pair-wised information, and has achieved great successes in discovering latent hierarchy in single-cell RNA-seq data^56^. Upon being mapped in Poincaré disk, radial coordinate of a node reflects its flow hierarchy, which describes its level in the latent information flow of network^55-57^. Representation of nodes by low-dimensional vectors can be employed in downstream analysis such as relationship mining or classification^58-60^. In this work, we first characterized chromatin structure with graph-theoretical statistics, discovering changes of network hierarchy in cancer and cell development processes. We then embedded chromatin contact networks into Poincaré disk using algorithms written by Nickel and Kiela^55^ to fulfill visualization and feature extraction purposes. The main purport of this work is schematically shown in Fig. 1. To the best of knowledge, this is the first study to implement Poincaré embedding to decipher structure of biomacromolecule aggregates with latent hyperbolic geometry.

**Fig. 1.**
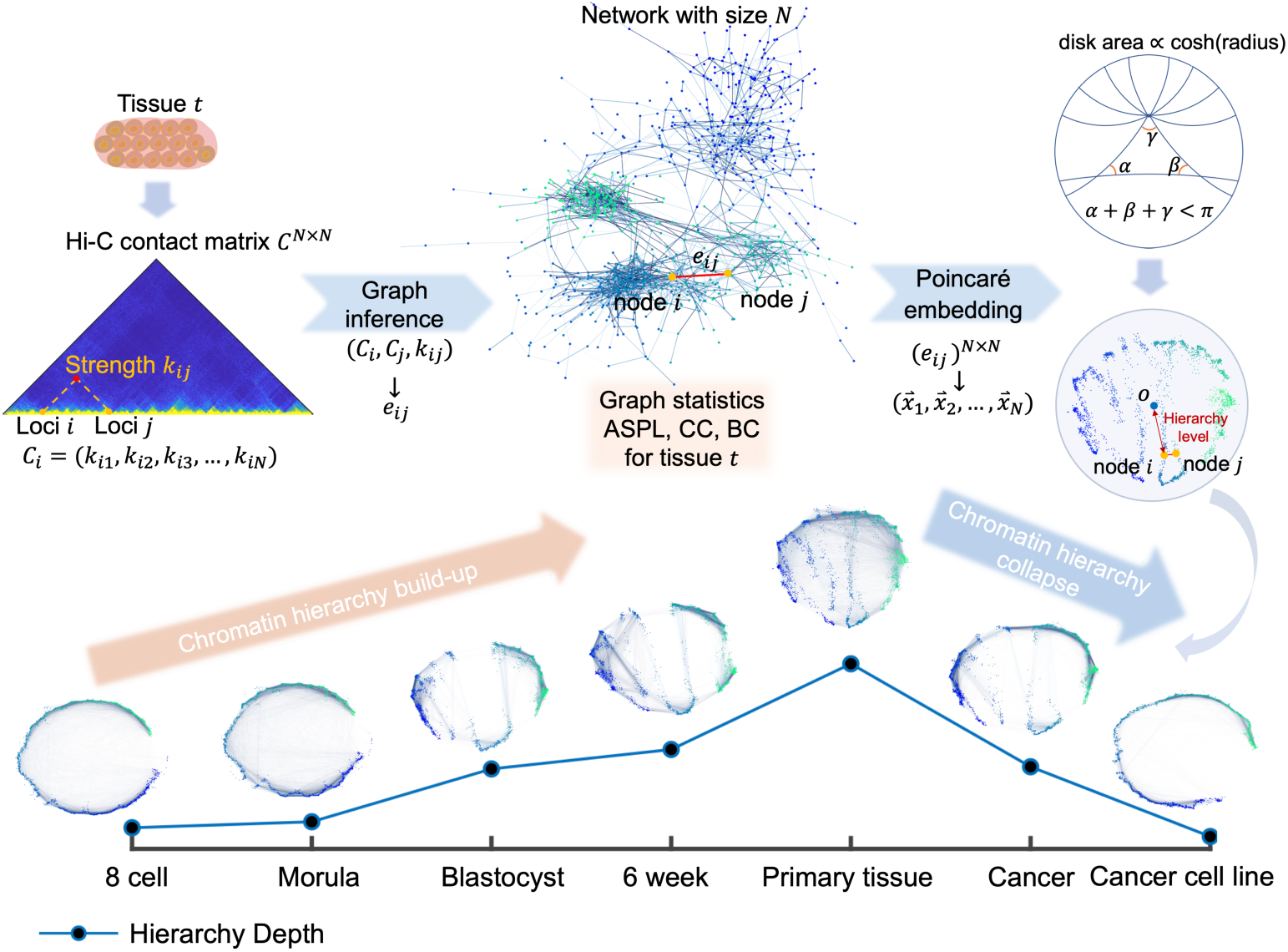
Graph statistics and Poincaré embedding reflect cell-state related changes in chromatin contact networks (CCNs). We performed network inferences on Hi-C datasets, followed by characterization of graph statistics including betweenness centrality (BC), average shortest path length (ASPL) and closeness centrality (CC), observing remarkable changes in network properties of CCNs in cancer and cell differentiation process. Then we embedded CCNs in Poincaré disk, which is a unit circle equipped with hyperbolic geometry, where area grows exponentially with radius. A hyperbolic triangle and hyperbolic lines were shown in schematic Poincaré disk. Through Poincaré embedding, hierarchy levels of nodes are manifested by radial distribution in disk. Hierarchy depth reflected by median Poincaré radii undergoes an initial increase during early embryo development, which continues in cell differentiation. In cancer, the hierarchy depth decreases and the trend continues from cancer to cancer cell lines.

## Result

### Graph statistics reflect changes of CCNs in cancer and cell differentiation

We examined changes in information transmission properties of CCNs from normal to cancer cells using published Hi-C datasets of CRC and T-ALL^32,35^. We first performed a graph inference to each intra-chromosomal Hi-C matrix (see in Methods). We then characterized the traffic fluxes of edges in CCNs using betweenness centrality^61^ (BC, see in Methods). BC of an edge measures its average probability to constitute the shortest path between any node pair, thus edges with high BC are essential for information transmission in a functional network^62^. The expected BC of edges at a sequential distance is positively related to the sequential distance it strides across, which underlines the bridge-like characteristics of long-range contacts (Fig. 2a). We next identified the normal- and cancer-specific edges in CRC, T-ALL and corresponding normal controls. For instance, an edge existing in more than 75% normal CCNs and concurrently absent in more than 75% cancer CCNs was defined as normal specific for a certain cell type. In both cases of CRC and T-ALL, we found significant decrease in linear sequential distances for cancer-specific edges compared with normal-specific edges (Fig. 2b, *p* = 1.6 × 10^−12^ for CRC and *p* = 2.7 × 10^−154^ for T-ALL). The analogous pattern of edge re-distribution shared by tumor and leukemia indicates a generality of cancers, which represents a loss of bridge-like contacts striding across long sequential distances.

**Fig. 2.**
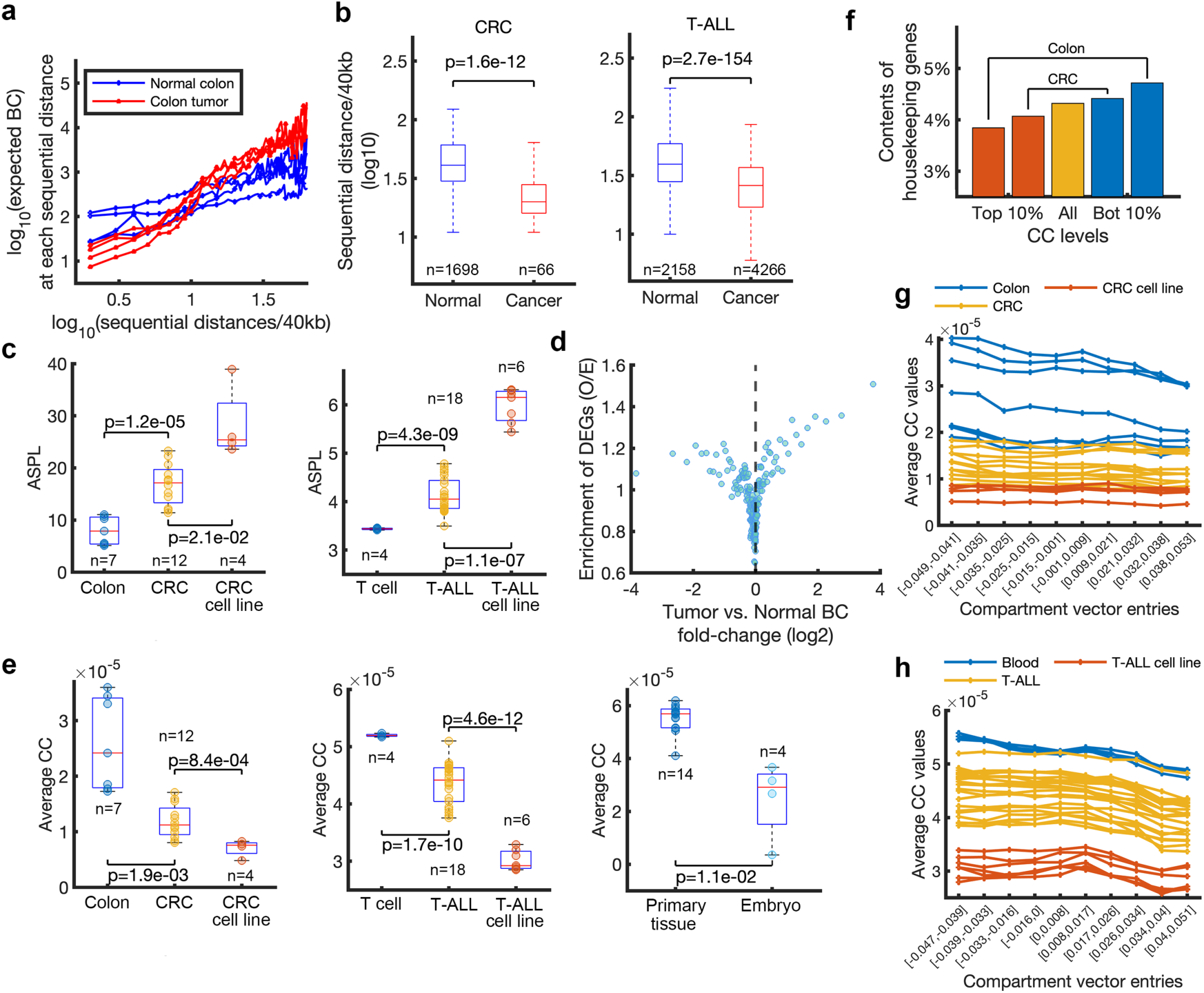
Decrease of information transmission efficiencies in cancer CCNs. **a** Logarithm plots of expected BC at various sequential distances for normal colon (blue) and colon tumor (red) samples from four individuals. **b** Boxplots showing logarithms of sequential distances of cancer or normal-specific edges in colorectal cancer (CRC) and T-lineage acute lymphoblastic leukemia (T-ALL). **c** Boxplots showing ASPL in cancer, cancer cell lines and corresponding normal controls for CRC (left) and T-ALL (right). **d** Differentially expressed gene (DEG) enrichment in edges with different levels of BC fold change (*FC*_*b*_). Edges were stratified into 200 groups according to *FC*_*b*_ rank. The x and y coordinates of points are averages of *FC*_*b*_ and DEG enrichments in CRC. **e** Boxplots showing average CC in cancer, cancer cell lines and corresponding normal controls for CRC (left) and T-ALL (middle). The right panel shows average CC of primary tissues and early embryos. **f** Bars showing contents of housekeeping genes (HKGs) in nodes with the top and bottom 10% CC levels in normal colon and CRC samples respectively. The average content of HKGs in the whole chromatin is also shown. **g-h** Plots showing relationships between CC values and compartment vector entries in CRC, T-ALL, their corresponding normal controls and cancer cell lines. The x and y coordinates of points show averages of compartment vector entries and these of CC values. Identification of normal and cancer specific edges and graph statistics including BC, ASPL and CC were carried out on intra-chromosomal CCN of chromosome 1. In each boxplot, the center line, the bottom hinge, the top hinge and the whiskers indicated the median, the 25^th^ percentile, the 75^th^ percentile and 1.5 times the interquartile range (IQR) of each dataset, respectively.

Considering the function of bridges for efficient routing in networks^15^, we anticipated that information transmission efficiencies of CCNs should decrease in cancer. We verified the conjecture by characterizing the average shortest path length^63^ (ASPL, see in Methods) of CCNs, the inverse of which measures the global information transmission efficiency of networks^64-66^. We found that cancer CCNs have significantly longer ASPL than normal CCNs in CRC and T-ALL (Fig. 2c, *p* = 1.2 × 10^−5^ for CRC and *p* = 4.3 × 10^−9^ for T-ALL), indicating a significant decrease of global information transmission efficiency in cancer CCNs. Interestingly, CCNs of cancer cell-lines have even longer ASPL than cancer CCNs, underlining distinctions between cancer and cancer cell lines (Fig. 2c, *p* = 2.1 × 10^−2^ for CRC and *p* = 1.1 × 10^−7^ for T-ALL).

We further investigated the possible influences of bridge-reorganization in CCNs in CRC. Edges were stratified into groups according to their BC fold changes (*FC*_*b*_) from normal colons to tumors. *FC*_*b*_ of an edge *e* is defined as the fold change of normalized BC,

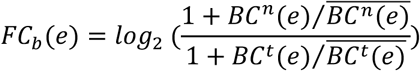

where *BC*^*n*^(*e)* and *BC*^*t*^(*e)* stand for BC of the edge *e* in a normal or tumor CCN respectively, and the overbar means averaging over all samples in the normal or tumor set. The normalization of BC was to filter noises and preserve large differences^67,68^. We inquired the TCGA database for gene expression profiles of normal colon and CRC samples and identified differentially expressed genes (DEGs). Notably, edges with high absolute *FC*_*b*_ tend to show enrichment in DEGs (Fig. 2d, which was defined as the observed/expected contents of DEGs in either end of edges (genes). For edges robustly changed in BC (absolute *FC*_*b*_ larger than 1), interestingly, their consisting nodes have around 80% intersection, indicating the occurrence of collective bridge-end switches in CCNs. Intersected bridge ends are enriched with genes involved in regulation of acute inflammatory responses and complement activation like C4BPA, CD46, PARK7 and SUSD4 (FDR<0.05, Supplementary table 3). Nodes on the other sides of broken bridges (FC_b_ < −1) in CRC include genes positively regulating stem cell proliferation (such as PTPRC, ASPM), whereas other ends of newly formed bridges (FC_b_ > 1) include genes regulating immunoreactive pathways (e.g., SLAMF1, MUC1, IRF6).

We speculated that nodes in CCNs undergo divergent changes in cancer. Aside from characterizing global routing efficiencies with ASPL, we further quantified routing efficiency of each node by closeness centralities^69^ (CCs) which measure the ASPL from one node to all other nodes (see in Methods). As expected, CC values are significantly decreased in CRC and T-ALL compared with their normal controls (Fig. 2e,*p* = 1.9 × 10^−3^ for CRC and *p* = 1.7 × 10^−10^ for T-ALL), and such a tendency continues from cancer to cancer cell-lines (*p* = 8.4 × 10^−10^ for CRC and *p* = 4.6 × 10^−12^ for T-ALL). Global loss of CC indicates a localization tendency of networks^70^. We identified nodes with the lowest 10% CC in normal colon to be more enriched with HKGs than average (Fig. 2f), corresponding to the fact that HKGs are often located at TAD borders^71,72^. The latter are known to form relatively insulated chromatin loci^73-75^. Nodes with the highest 10% CC in normal colon are less likely to contain HKGs than average (Fig. 2f). They instead involve genes regulating chromatin silencing (e.g., DNMT1, HIST1H2AB, and RIF1) and chromatin structure stability (e.g., CENPQ, NAP1L1, and DCP2). We normalized CC along the sequence into its Z-score vector and identified nodes with significant changes (p < 0.05) in CRC, among them are genes involved in regulation of cell adhesion (e.g., PCDH1, DSG2, and FAT2), negative regulation of immune response (such as IL10, CD34, BAP1, and CD6), regulation of extracellular matrix assembly (e.g., AGT, HAS2) and regulation of cell proliferation (e.g., MYC).

For each Hi-C sample, we stratified chromatin loci into groups according to their corresponding entries in the compartment vector. Interestingly, a negative correlation was observed between CC and compartment vector entries in normal samples, which declined in CRC and T-ALL and indicated a loss of node heterogeneity in cancer (Figs. 2g, h). While decreases of CC are occurring genome wide, the most compartment-B nodes suffer from more severe decreases in colon tumors (Supplementary Fig. 1a). The prominent susceptivity of these most compartment-B like or compact nodes to cancer reflected by severe decrease of CC is reminiscent of reported progressive heterochromatin decompaction in tumors^31^.

Intrigued by the stemness in cancer cells^37,38,40-42^, we also looked into the cell differentiation process, during which chromatin structure undergoes significant changes^45,76^. Here, we studied changes in CCNs in differentiation using published Hi-C datasets of human early embryonic cells and human primary tissues^77,78^. Early embryos from 2-cell to blastocyst possess omnipotency, while primary tissues are terminally differentiated. In the set of primary tissues, significantly higher CC values were observed relative to the set of early embryos (Fig. 2e, right; *p* = 1.1 × 10^−2^). The changes observed in graph statistics indicate an increase in routing efficiency of CCNs in cell differentiation, opposite to what happens in cancer. Furthermore, compared with primary tissues, stem cells resemble tumor cells in that their most compartment B-like nodes also have low CC values (Supplementary Fig. 1c). Interestingly, heterochromatin in stem cells shows structural openness^79^ that is analogous to the heterochromatin decompaction in cancer^31^.

### Poincaré embedding of CCNs reveals cell-state related changes in hierarchy depths

To visualize the structure of chromatin contact networks, we next embedded Hi-C of each sample in the Poincaré disk. Graph embedding is a dimensionality reduction strategy which transforms graph connectivity into distances in the embedding space^52,53^. It is known that the existence of hierarchically nested TADs in Hi-C matrix leads to strong clustering in the contact patterns^80,81^, and at the same time, power-law degree distributions of chromatin contact networks indicate the existence of node heterogeneity^12^. Strong clustering and node heterogeneity characterize a network with intrinsic flow hierarchy^15^. As discussed above, Poincaré disk is compatible with the underlying hyperbolic geometry in flow-hierarchical networks^15^, like CCNs. Through Poincaré embedding, nodes with higher hierarchy statuses are scattered around the disk center, while nodes with lower hierarchy statuses are distributed near the disk periphery^15,55^. Specifically, we embedded each intra-chromosomal CCN into Poincaré disk at 40kb resolution adapting the algorithms of Ref. ^55^ (see in Methods).

The simple two-dimensional coordinates of mapped chromatin loci can be used to describe concisely the chromatin properties, among many of which, chromatin compactness is important for its indication of a silent transcription environment^82,83^. To quantify the local compactness of embedded chromatin structure, we defined the metric neighborhood radius (*N*_*r*_ see in Methods). For each bin, its *N*_*r*_ is the average Poincaré distance between all pairs of nodes within its neighborhood, which then describes how compact the local environment is. As shown in Fig. 3a, local maxima of *N*_*r*_ are located at TAD boundaries and contain 46% more HKGs than the average. Loci of the lowest quartile of *N*_*r*_ are located at TAD interior, with HKGs 34% more depleted than the average. These low-*N*_*r*_ loci include genes with functions of chromosome structure stability maintenance (e.g., CENPO, HAT1, and HJURP). These observations suggest that nodes within the same TAD share similar contacting profiles, thus appear as a network community and form a cluster in Poincaré disk, while TAD boundaries are manifested as cluster borders.

**Fig. 3.**
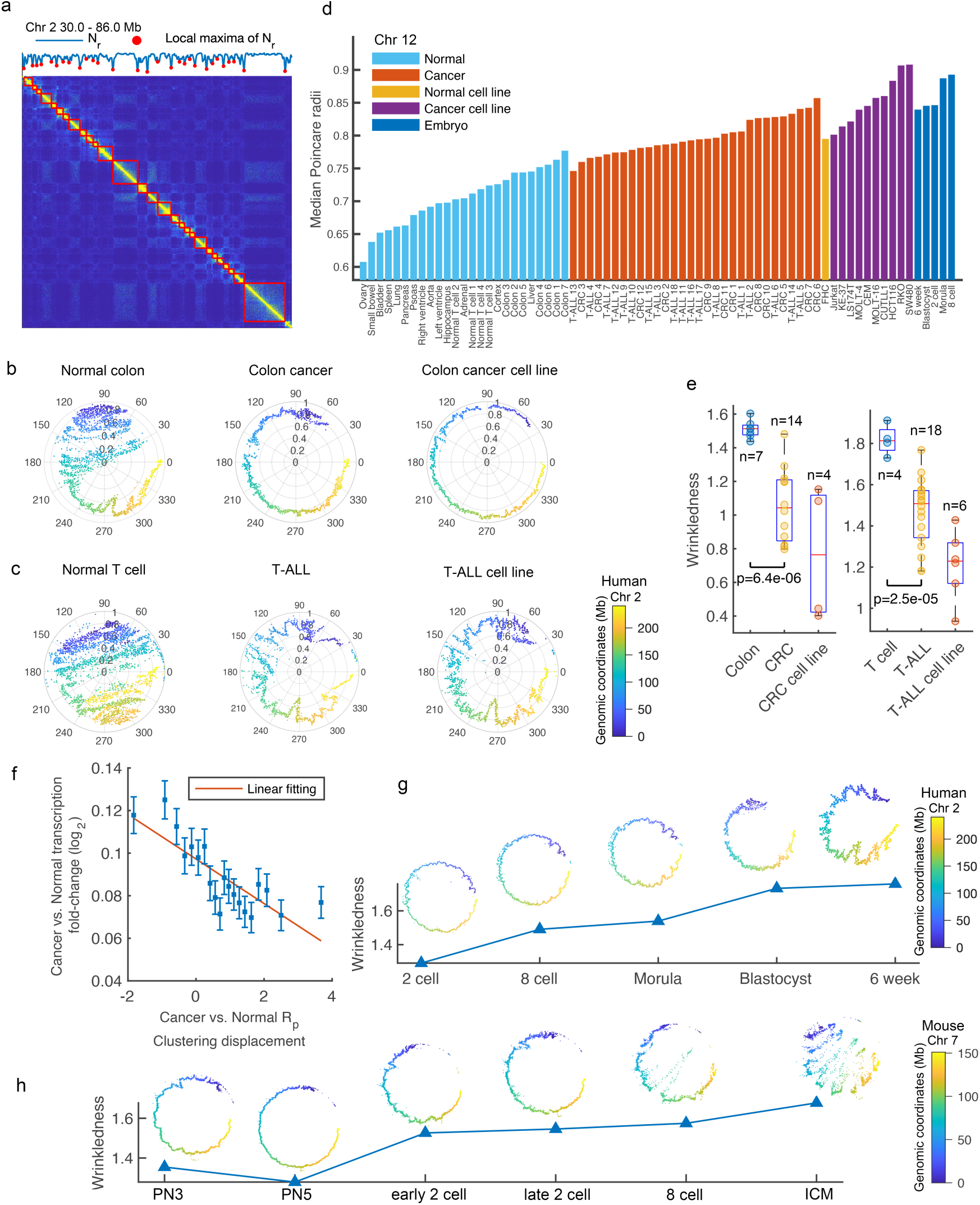
Poincaré embedding of CCNs shows cell-state related hierarchy changes in cancer and early embryo developments. **a** Correspondance of local maxima of neighborhood radius (*N*, see in Methods) and TAD boundaries in a normal T-cell Hi-C. **b-c** Embedded structures of CCNs in Poincaré disk for normal controls (left), cancer (middle) and cancer cell lines (right) of CRC and T-ALL. The color distribution of embedded nodes represents DNA sequential order. **d** Bars showing median Poincaré radii (*R*_*p*_) of each sample with cell types varying from primary tissues, cancer, cancer cell lines and early embryos. **e** Boxplots showing wrinkledness (*W*_*k*_, see in Methods) in cancer, cancer cell lines and corresponding normal controls for CRC (left) and T-ALL (right). **f** Scatter plot showing relationship between cluster displacements (see in Methods) of *R*_*p*_ and transcripts per million (TPM) fold changes from normal colon set to colon tumor set. All genes were stratified into groups according to cluster displacements of *R*_*p*_ ascendingly. The x and y coordinates of points are group averages of cluster displacements of *R*_*p*_ and that of TPM fold changes. The lower and upper bounds of error bars indicate 95% confidence intervals of TPM fold changes. Residue of linear fitting is 0.6504. Embedded structures of CCNs and corresponding changes in *W*_*k*_ during early embryo developments of human (**g**) and mouse (**h**).

We embedded individual intra-chromosomal CCNs of each sample into Poincaré disk, considering the existence of chromosome territories. Discriminations between normal and cancer chromatin structures through Poincaré embedding are clearly seen in Figs. 3b, c, for both CRC and T-ALL. Changes in cancer manifest as two main features: the shift of the structure from the center of disk to the periphery, and the decline of the structural “wrinkledness” (*W*_*k*_, see in Methods). Similar tendencies from normal to cancer were observed in CRC and T-ALL, and continue in the change from cancer to cancer cell lines (Figs. 3b, c).

In term of radial distribution, almost all chromosomes show significant changes in that embedded structures relax towards the disk periphery in CRC and T-ALL compared to their corresponding normal controls (Supplementary Fig. 2a). Increases of Poincaré radii (*R*_*p*_) reflect the reduction of flow hierarchy depth of CCNs in cancer, since the nodes originally located near the disk center in normal cells lose their advantageous positions to influence the whole network in cancer cells. In CRC, nodes with their Poincaré radii (*R*_*p*_) significantly increased (*p* < 0.001) consist of genes regulating chemokine and interferon secretion (GATA3, CD2, IL33 etc.) and negatively regulating MAPK activity (PTPN6, CBLC, IL1B etc.), such genes undergo most severe hierarchy decrease in the contact network. Generally, *R*_*p*_ shows an ascending trend from normal tissues to cancer, as well as from cancer to cancer cell lines (Figs. 3b, c, d). Interestingly, as shown in Fig. 3d, median *R*_*p*_ of normal colon samples from adjacent normal areas in tumor resection specimens are significantly higher than those of primary tissues (*p* = 2.1 × 10^−7^) but lower than those of cancer samples (*p* = 1.7 × 10^−5^), indicating the possible distinguishment between tissues at risk for tumorigenesis and healthy tissues in chromatin structure. Without data on normal colon tissue from non-cancer donors, we compared the embedded chromatin structures of normal colons adjacent to tumor and that of small bowel. While *R*_*p*_ of colons adjacent to tumor is higher than that of small bowel, we identified nodes with *R*_*p*_ robustly higher in the former set. Specifically, we transformed the *R*_*p*_ vector of small bowel into a Z-score vector by subtracting the average and dividing by the standard deviation of *R*_*p*_ vectors of colons adjacent to tumor. We defined nodes with Z-score smaller than -10 as showing robustly higher *R*_*p*_ in colons adjacent to tumor than in small bowel. These nodes contain genes with enriched pathway of positively regulating neutrophil chemotaxis (FDR<0.005, Supplementary table 4), including CCL19, CXCL1, DAPK2, PPBP. Moreover, *R*_*p*_ level of normal colon cell line is between those of normal colons and CRC cell lines (Fig. 3d), probably implying a malignancy related abnormality in chromatin structures of normal cell lines.

Direct visualization of embedded structures implies the loss of chromosome fine structure features in cancer. To quantify the wrinkledness of embedded structure, we defined the descriptor *W*_*k*_ based on structure curvature calculation (see in Methods). *W*_*k*_ of cancer samples are significantly lower than that of their normal counterparts at the mega-base scale, while the structures of cancer cell lines are even less wrinkled than corresponding cancer samples (Fig. 3e). The increase in *R*_*p*_ reflects a decay in global hierarchy depth, while the decrease in *W*_*k*_ reflects loss of node heterogeneity. Analogous changes occur on all individual chromosomes(Supplementary Fig. 2b, indicating that the decline of CCN hierarchy depth occurring genome-widely may be a generality for cancer.

The role of chromatin structure in gene expression regulation has raised great interest and remains relatively poorly understood^84-87^. To investigate the correlation between changes of chromatin structure and that of transcription in cancer, we further used the published transcription profiles on the normal colon and colon tumor tissues^32^. Fold change of transcripts per million (TPM) was used to measure expression change for each transcript, while the cluster displacement (*d*_*U,V*_, see in Methods) of *R*_*p*_ was calculated to quantify the hierarchy status change of each node in CCN. We calculated the *d*_*U,V*_ of each node’s *R*_*p*_ from the normal colon cohort *U* (n=7) to the colon tumor cohort *V* (n=12). Notably, we observed a negative correlation in collective behavior of TPM fold changes and *d*_*U,V*_ (Fig. 3f; Pearson’s r = -0.806, *p* = 1.7 × 10^−5^), indicating a tendency that genes with increasing hierarchy statuses in contact networks are transcribed more frequently changing from normal to tumor. The correlation between changes in CCN and that of transcription implies that certain genes have their roles shifted in both repertoires of chromatin structure and expression. Among nodes acquiring the largest decrease in hierarchy (top 5% of *d*_*U,V*_ are genes regulating cell migration (EGFR, NRP1, and CXCL12, etc.) and regulating chemokine and interferon secretion (IL33, CD244, and C5, etc.), while nodes undergoing the most increase in hierarchy (bottom 5% of *d*_*U,V*_) contain genes regulating antigen processing and presentation via MHC class II, such as KIF5A, HLA-DMA, and DCTN2, and those regulating Wnt pathway, such as WNT11, CTNNB1, and PPP3CB, among others.

We also investigated the early embryonic and cell differentiation processes by Poincaré embedding to associate with the de-differentiation traces in cancer cells. We first studied changes in CCNs during early embryonic development using published Hi-C data on human and mice^44,77^. Compared to carcinogenesis, CCNs change in an opposite way during early embryonic developments. Build-up of genome architecture before ICM (inner cell mass) is manifested by an increased *W*_*k*_ in embedded structures (Figs. 3g, h and Supplementary Figs. 2c, d). The increase of *W*_*k*_ from 2-cell embryo to blastocyst or ICM is shared by both human and mice. In embryonic development of mice, increasing trends of *W*_*k*_ were observed in both paternal and maternal alleles (Supplementary Fig. 2e). Early embryo development initiates the cell differentiation process, and primary tissues are at the other end of cell differentiation trajectory^88,89^. We investigated changes in CCNs in cell differentiation by comparing the human early embryo (from 2-cell to blastocyst) and primary tissue samples using published Hi-C datasets^77,78^. From early embryos to differentiated tissues, remarkable decrease of *R*_*p*_ and increase of *W*_*k*_ were observed genome-widely (Supplementary Figs. 2a, b), reflecting the establishment of CCN hierarchy in cell differentiation.

In summary, CCN undergoes an initial establishment of hierarchical architecture during early embryo development followed by further development into a more complex hierarchical structure in cell differentiation. In cancer, however, CCN collapses into a less structured organization largely dictated by the 1-D linear DNA sequence.

### Hierarchy disorder of CCNs in cancer

From foregoing analysis, the hierarchy depth is an important descriptor of a chromatin contact network. On the other side, we anticipated that hierarchy order of nodes in CCN is also vital and related to cell states. We then quantified flow-hierarchy statuses of individual nodes in each chromosome using Poincaré radii (*R*_*p*_) obtained from embedded intra-chromosomal CCNs, since nodes with smaller *R*_*p*_ are closer to all nodes in the whole network and have more advantageous positions in initiating the information spreading^15,55,90,91^.

To verify the ability of flow-hierarchy statuses of nodes to characterize the cell states, we transformed the *R*_*p*_ vector into its Z-score vector along sequence. Such an operation was performed to evaluate each node’s relative flow-hierarchy status among all nodes. We next performed a principal component analysis (PCA) on *R*_*p*_ vectors for all samples, based on which each sample was represented by a low-dimensional vector in the principal component (PC) space. In both cases of CRC and T-ALL, samples are departed in PC1 according to their belonging clusters of normal, cancer and cancer cell line (Fig. 4a). The fact that malignancy states of Hi-C samples can be distinguished by relative hierarchy of nodes inspired us to further study the association between cell identity and hierarchy rank of nodes. We then evaluated the similarity in hierarchy rank using the Spearman’s correlation coefficient between *R*_*p*_ vectors of different samples. Conservation of intra-chromosomal hierarchy order among normal colons was observed, whereas weakened among colon tumors (Fig. 4b). Notably, robust distinctions in hierarchy order between normal colon and colon tumor were shown, indicating the occurrence of hierarchy disorder in CCNs in cancer (Figs. 3b, c). On the other hand, each normal primary tissue has a unique hierarchy order, except for two pairs (cortex and hippocampus, left ventricle and right ventricle), which have high (larger than 0.9) Spearman’s coefficients (Fig. 4c). This result further underlines that hierarchy order of CCN not only reflects distinguishment between normal and cancer tissues, it is a delicate description of normal tissue specificity.

**Fig. 4.**
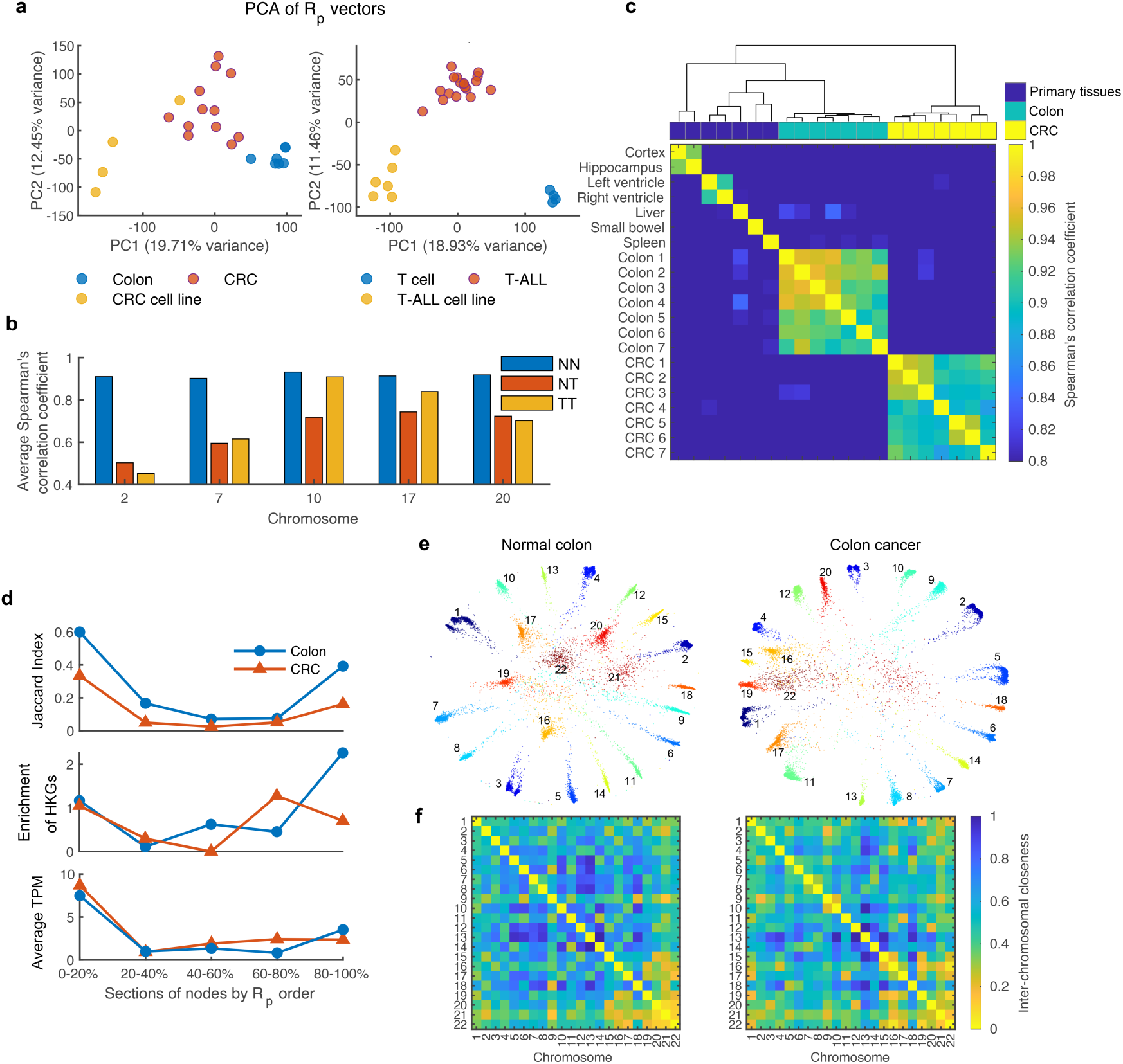
Hierarchy disorder of CCNs in cancer. **a** PCA for *R*_*p*_ vectors of samples in CRC and T-ALL datasets. Each point represents a sample in PC space defined by the first 2 principal components. Normal, cancer and cancer cell lines were discriminated by PC1 in both CRC and T-ALL. **b** Bar graphs showing average Spearman’s correlation coefficients of *R*_*p*_ between all pairs of normal colons (NN), between all pairs of colon tumors (TT) and between each normal colon and each colon tumor (NT). **c** Rank similarity in *R*_*p*_ among samples including primary tissues, normal colons and colon tumors. Each unit color in the heatmap represents the value of Spearman’s correlation coefficient between *R*_*p*_ vectors of a sample pair. Samples were ordered according to Ward’s hierarchical clustering method. **d** Jaccard index (top), enrichment of HKGs (middle), and average TPM of genes (bottom) of each hierarchy class for the normal colon and CRC set. Enrichment of HKGs was calculated as the observed/expected proportion. Statistics in **c** and **d** were from intra-chromosomal CCNs of chromosome 10, therein normal colon and CRC sets both achieved fine conservation of hierarchy order (**b**). **e-f** Embedded structures of whole chromatin contact networks and inter-chromosomal closeness matrices for a pair of normal colon (left) and colon tumor (right) from one individual. Each heatmap entry ranging from 0 to 1 indicates the closeness between each pair of chromosomes.

We inquired the stability of hierarchy order by calculating the Jaccard Index (JI) for each 20% section of nodes in hierarchy order. Specifically, for each sample, nodes in an intra-chromosomal CCN were stratified into five classes according to its entry in the *R*_*p*_ vector (i.e., sections of 1^st^ to 20^th^, 20^th^ to 40^th^, …, 80^th^ to 100^th^ quantiles). For the normal colon and the colon tumor sets (n=7 for both), the size of intersection is divided by the size of union, hence yielding the JI for each of the five hierarchy classes. The class with the highest hierarchy has the largest JI in both normal and tumor sets (Fig. 4d), indicating strong conservation in node composition of corresponding class. Interestingly, JI for the class with the lowest hierarchy shows similar conservation, while node compositions of middle classes are less conserved. In each class, node composition for the normal set is more conserved than that for the tumor set, reflecting the higher heterogeneity of cancer than normal cells.

In normal colons, as seen in Fig. 4d, the class with the lowest hierarchy is enriched with housekeeping genes (HKGs). We reason that HKGs are often located at insulated TAD boundaries where chromatin loci have localized contacting profiles^71,72^, which correspond to distal parts of CCNs hence have low flow-hierarchy. In colon tumors, however, enrichment of HKGs in the lowest-hierarchy class weakens (Fig. 4d), which is consistent with the disturbed insulation at TAD boundaries in cancer^32^. We calculated average transcripts per million (TPM) of genes in each class for normal colon and colon tumor sets using published transcription profiles^32^. Notably, coexistence of high gene transcription level and strong conservation (depicted by JI) in the highest -hierarchy class were observed in both normal colon and colon tumor sets (Fig. 4d). We derived the difference set for genes in the highest-hierarchy class for either sample set, that is, genes which are in the highest-hierarchy class of normal set but down-graded in tumor set and vice versa. Among the normal-specific genes with the highest hierarchy in CCNs, PTEN, RET, CXCL12, and RBP4 are involved in negative regulation of cell proliferation, lymphocyte migration and maintenance of gastrointestinal epithelium, while TET1, DLG5, and DNA2 in the cancer-specific set involve in embryonic stem cell maintenance, epithelial-mesenchymal transition (EMT) and DNA replication checkpoint. Correspondingly, loss of PTEN expression is reported in 34.5% of CRC resultant from generic or epigenetic mechanisms^92,93^.

The foregoing analysis thus revealed severe hierarchy collapse and disorder in intra-chromosomal CCNs in cancer. We further investigated changes in the context of whole chromatin during the same process. Due to existence of chromatin territories, intra-chromosomal contact networks are relatively independent components of the whole-chromatin CCN. We embedded the whole-chromatin CCNs in Poincaré disk at 100kb resolution following a procedure analogous to aforementioned intra-chromosomal CCN embedding (see in Methods). As shown in Fig. 4e, chromosomes 16,17,19,20,21 and 22 are located near the disk center for normal colon, whereas relocate towards the disk periphery in colon tumor. We reasoned that chromosomes possess unequal hierarchy statuses owing to inhomogeneous radial distribution in nucleus. In fact, short and gene-rich chromosomes 16,17,19,20,21 and 22 are reported to be located in nucleus interior ^94,95^, and have wide contacts with other chromosomes^5^. We defined the inter-chromosomal closeness matrix (see in Methods) to evaluate the distance between each pair of chromosomes relative to corresponding intra-chromosomal distances, so that the matrix entry ranging from 0 to 1 represents the closeness between each pair of chromosomes. As shown in the closeness matrices for the normal colon CCN, long chromosomes are far from each other while short chromosomes are close to each other and not far away from long chromosomes, except for chromosome 18 which has a behavior similar to long chromosomes (Fig. 4f). However, for colon tumor CCNs, the distinct contacting patterns of short and long chromosomes are attenuated, indicating disturbed segregation of long chromosomes. Such a change corresponds to findings that long chromosomes are enriched with lamina associated domains (LADs)^96^ and that lamina associated heterochromatin are relocated into the nuclear interior during tumorigenesis^32^.

## Discussion

### Decrease of hierarchy depth in CCNs in cancer hinders the efficient genetic information transmission

The frequently mentioned hierarchy in chromatin in previous studies describes nested hierarchy, while in this study we investigated the flow hierarchy of networks, which describes the influences of nodes in the latent information flow. The hierarchy of CCNs characterized with graph statistics and graph learning techniques both uncovered a significant decrease in hierarchy depths of CCNs in cancer. Network hierarchy depth is an important property, it determines how stable the network is against perturbance^97^. Nodes with high hierarchy statuses are vital in maintaining network performance and stability^97-99^. We first applied graph statistics to CCNs, including centralities and path lengths which are correlated with flow-hierarchy depth^11,15,91^. We then performed graph embedding and obtained another flow-hierarchy descriptor *R*_*p*_ based on a machine-learning method^55^. With both analytical and inferred metrics for network hierarchy, we observed consistent decrease in hierarchy depth and hierarchy disorder of CCNs in cancer.

The main benefit for entities to be organized into a network is to serve the aim of information transmission. Hierarchical networks are well known for their efficient routing guaranteed by the underlining hyperbolic geometry^15^. Chromatin contact network is a complex hierarchical network characterized by the scale-free degree distribution and existence of nested TADs^12^. In a chromatin contact network, where a physical contact can be random or bear the mission of transmitting genetic signals (e.g., enhancer-promoter interactions), long-range interactions serve as bridges and can increase the genetic signal transmission efficiency^15^. In different cancers, breaking of these bridges (loss of long-range contacts in Hi-C) is expected to hinder the global information communication, manifested by decrease of CC as well as the increase of ASPL.

While chromatin is a three-dimensional Euclidean object, hierarchical chromatin contacting networks have intrinsically a hyperbolic geometry, thus hyperbolic representations can reserve chromatin structural information well. We performed Poincaré embedding of CCNs, which allows easy visualization of subtle structural properties revealed by Hi-C. Then we used *R*_*p*_ to measure hierarchy statuses of nodes: a node with smaller *R*_*p*_ is closer to all other nodes in network and is more advantageous in initiating information spreading^91,100^. Normal tissues are characterized with low *R*_*p*_ levels, indicating the existence of deep hierarchy depth in normal CCNs. In cancer, decrease of CCN hierarchy depth is reflected by significant attenuation of *R*_*p*_. Absence of nodes with small *R*_*p*_ can severely hinder efficient global routing of networks, leading to the decline of information transmission efficiency^15^. The fact that *R*_*p*_ levels of normal colon tissues adjacent to tumor are higher than those of normal primary tissues suggests that tissues at risk for cancer have characteristics between normal and cancer tissues. The other possibility is that these normal colon samples are slightly contaminated by cancerous cells. Further studies are needed to rule out the latter possibility. Another interesting discovery is that normal colon cell-lines are between normal colons and CRC cell lines in *R*_*p*_ level. We conclude that structural abnormality exists in chromatin of normal cell-lines, which may be correlated with their immortality. Sensitivity of CCN to tumorigenesis suggests its potential usage in cancer risk stratifications and early detection.

Aside from decrease of network hierarchy depth, the interior hierarchy order becomes less uniform in tumorigenesis. It is likely that individual nodes in CCN can transmit information through contacting with other nodes, the local or global contacting patterns of which are closely associated with the hierarchy ranks. The proper matching of nodes with their hierarchy ranks may be vital for accurate signal transmission, while disruption of which may result in dysregulation of network functionality. In CRC, node hierarchy orders by *R*_*p*_ are conserved among normal samples while reshuffled into more diverged states in cancer samples. Notably, with both metrics of CC and *R*_*p*_, we observed significant changes of hierarchy statuses on a set of genes regulating chemokine and interferon secretion and embryonic stem cell maintenance, indicating existence of anomalous immune responses and embryonic stem cell-specific signatures in cancer^101-104^. The observed hierarchy disorder of CCNs in cancer resembles a transition of glassy states in hierarchical networks under disturbances, consistent with the deep-to-shallow hierarchy transition^105^. The cancer-specific hierarchy order of chromatin loci is worth attention, since the generality emerging in the torrent of tumor heterogeneity may be a prerequisite for cancer cell survival.

### Evolvements of CCNs in cancer, early embryo development and cell differentiation

This study shows that Poincaré embedding can faithfully and sensitively reveal variations in chromatin structure especially in processes during which the CCN hierarchy depth significantly changes, including cancerization and cell developments. From results of *R*_*p*_ levels, cancer and cell differentiation are accompanied by decrease and increase of hierarchy depth, respectively. During early embryo development of human and mice, hierarchy depth of CCNs gradually increases. Interestingly, as shown in Supplementary Fig. 2d, embedded chromatin structure is more quasi-1D-like at PN5 and late-two-cell stages in mice, reflected by a drop of wrinkledness (*W*_*k*_). Such variations in chromatin structure coincide with minor and major ZGA events, during which large-scale gene transcription starts^106,107^. Therefore, increased smoothness indicated by *W*_*k*_ decrease in chromatin scaffold appears to occur with transcription machinery assembling, since RNA polymerase II (Pol II) needs to be loaded onto genome during minor ZGA and extended to whole genome during major ZGA^108^. Comparing CCNs of blastocyst and primary tissues, we found that the hierarchical CCN of blastocyst embryo is initially built up but only become completely established upon further differentiation into downstream monofunctional cells.

In cancer, hierarchical CCNs collapse to resemble stem cells, reflected by both metrics of CC and *R*_*p*_. In term of *R*_*p*_ levels, cancer tissues are between normal tissues and early embryos, implying the stemness in cancer chromatin structures. Cancer tissues resemble pluripotent embryos also in their significantly lower CC values in CCNs compared with normal tissues. Notably, in normal tissues, heterochromatin nodes with the lowest compartment vector values have the highest CC. These heterochromatin nodes with the putative most compact environment undergo the most severe CC reduction in tumor, corresponding to evident decompaction of heterochromatin observed during tumor progression^31^. Since nodes possessing high CC are vital for network stabilities^97^, we believe that further study should be carried on these heterochromatic loci which may be potential targets for cancer diagnosis and treatment. Negative correlations between CC and compartment vector were observed in primary tissues, but not in cancer. Existence of such negative correlations in normal tissues reflects the compact nature of heterochromatin. Heterochromatin loci separated by long sequential distances are packed into compact clusters and have strong physical associations therein^109,110^. Long-range contacts among heterochromatic loci act as bridges in CCN, contribute to the high CC of heterochromatin nodes. We assume that progressive heterochromatin decompaction in cancer^31^ narrows the gap in compactness between heterochromatin and euchromatin, thus weakens the heterogeneity of CC.

The observation that CCNs in both cancer and pluripotent embryos have low hierarchy depth may originate from the correlation between network stability and hierarchy. The more hierarchical a network is, the more resistance and resilience it has against disturbance^97,98^. However, a network with deep hierarchy depth can jump to a state with shallow hierarchy depth under strong disturbance^105^. Since network instability leads to plasticity^111,112^, we speculate that the shallow hierarchy or quasi-1D-like characteristics in CCNs of embryos are related to their proficiency to differentiate into downstream variants. When it comes to cancer tissues, structural instability of CCNs may favor uncontrolled proliferated cells. However, chromatin structures of cancer cells and embryo stem cells are distinguished in the existence of a determined and programmed transition from shallow-hierarchy to deep-hierarchy statuses. Based on such an argument, we postulate that epigenetic modifications play important roles in determining the cell fate by monitoring node attributes in CCNs, i.e., consistent with the ability of aberrant epigenetic modifications to transform stem cells into cancer cells^43,77,113^. Based on partial similarity in chromatin structure and substantial differences in cell nature between cancer and stem cells, it would be interesting to understand their genetic differences to minimize the effects on stem cells in gene therapy for cancer.

## Methods

### Normalization of Hi-C and compartment assignment

For each sample, normalization was performed on the raw Hi-C matrix using the Iterative Correction and Eigenvector decomposition (ICE) method^114^, yielding the normalized Hi-C matrix. The normalization was performed on intra-chromosomal Hi-C matrices with 40kb resolution and the whole-chromatin Hi-C matrices with 100kb resolution. We used the standard PCA-based method^5^ to assign compartments on each chromosome for each Hi-C sample. Specifically, entries in a normalized Hi-C matrix were divided by the average values of entries at corresponding sequential distances, yielding an observed over expected (O/E) matrix. A correlation matrix was then derived by calculating the pairwise correlation coefficients of rows in the O/E matrix, followed by a PCA procedure on the correlation matrix. The sign of the PC1 vector combined with the GC content distribution was then used to determine the assignment of compartments.

### Closeness centralities, average shortest path length and betweenness centralities of chromatin contact networks

Similarity of node attributes and neighboring profiles has been widely used in predicting missing links in networks^115,116^. In our network inference procedure, we calculated the correlation matrix by computing the Pearson’s correlation coefficient between rows in the ICE-normalized Hi-C matrix. Pearson’s correlation method was used in inferring effective connectivity strengths between nodes in biological networks^117,118^, but to the best of our knowledge, it has not been applied to infer CCNs. We assigned the top 5% edges with the highest Pearson’s correlation coefficients as the connected links. In this way, an unweighted network was derived and fed into followed calculations of network metrics including closeness centrality (CC), average shortest path length (ASPL) and betweenness centrality (BC).

CC was calculated as the inverse sum of the shortest paths from a node to all other nodes in the graph^69^,

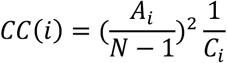

Where ***A***_***i***_ is the number of reachable nodes for node *i* (excluding *i*), ***N*** is the number of nodes in *G*, and ***C***_***i***_ is the sum of shortest paths from node *i* to all reachable nodes. Shortest paths of all node pairs were calculated with breadth-first search algorithm on unweighted graph^63^.

The betweenness centrality (BC) of an edge *e* is the sum of the fraction of all pair-wise shortest paths that pass through *e*^61^:

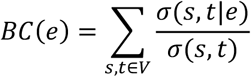

where *V* is the set of nodes, σ(*s, t*) is the number of shortest (*s, t*)-paths, and σ(*s, t*|*e*) is the number of these paths passing through *e*.

### TCGA gene expression analysis

In this paper, protein coding genes were downloaded from GENECODE^119^ and the reference genomes for human and mouse are hg19 and mm9, respectively.

We downloaded all available RNA-seq data of CRC and normal colons from TCGA database, which were then converted into TPM format. TPM fold change of each gene from normal colons to colon tumors was calculated as follows,

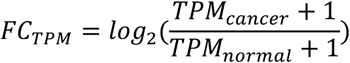

TPM_cancer_ and TPM_normal_ of a gene were calculated by averaging TPM in all cancer and normal samples respectively. We performed differential gene expression analysis with the R/Bioconductor package “DESeq2” ^120^. A significant expression change was defined by an absolute *FC*_*TPM*_ larger than 1 and a p-value lower than 0.05.

### Gene function analysis

In this study, Gene Ontology (GO) enrichment analysis of interested genes were performed using the R package ClusterProfiler^121^.

### Embedding chromatin contact networks in Poincaré disk

To embed intra-chromosomal CCNs, ICE-normalized Hi-C matrices of 40kb resolution were used as adjacency matrices. We used a Monte Carlo strategy to embed the weighted networks into Poincaré disk. First, we transformed a Hi-C matrix into a probability matrix by re-assigning values of the largest 25% elements as 1s and divided the rest elements by the 75^th^ quantile of Hi-C matrix. We then sampled an unweighted adjacency matrix based on the probability matrix at each training epoch. In embedding CCNs of the whole chromatin, we used ICE-normalized Hi-C matrices of 100kb resolution. The whole Hi-C matrix was transformed into an unweighted adjacency matrix by re-assigning entries ranging from the 85^th^ to 95^th^ quantile of whole matrix into 1s and the rest into 0s. The same adjacency matrix was used through all training epochs until convergence.

The unweighted graph at each epoch was embedded in Poincaré disk using algorithms written by Nickel and Kiela^55^. Loss function of each epoch was optimized by encouraging connected nodes in the unweighted graph to be close in Poincaré disk:

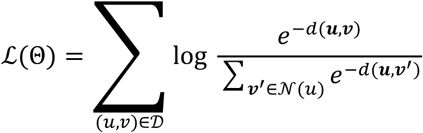

where 𝒩(*u*) = {ν | (*u*, ν) ∉ 𝒟} ∪ {*u*} is the set of negative examples, or, unconnected nodes for *u*. In Poincaré disk, hyperbolic distance was calculated as follows,

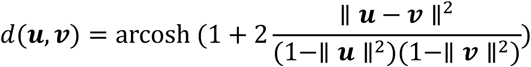

where ***u, ν*** are the embedded polar coordinates of node *u, ν*. The number of training epochs needed depends on convergence of loss function. The same training protocol was applied to all individual Hi-C samples and reproducibility of embedding was examined.

### Neighborhood radius

We defined the neighborhood radius *N*_*r*_ to describe local compactness of an embedded structure. Firstly, we obtained a distance matrix *D*_*p*_ by calculating Poincaré distances for all node pairs. A square window with length of 2*l* + 1 was then shifted along the diagonal of *D*_*p*_. At each shifting step, all matrix elements in the square were averaged and the average was defined as *N*_*r*_ of the bin at the square center.

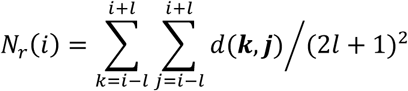

where *d*(***k, j***) is the hyperbolic distance between the *k*th and the *j*th node in Poincaré disk, *l* is the neighborhood window size. We used *l* = 5 for all calculations of *N*_*r*_.

### Wrinkledness of embedded structures

To describe the wrinkledness of embedded structure geometry, we defined *W*_*k*_ by calculating the mean value of local curvatures approximated by finite interpolation method:

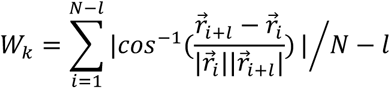

where *l* is the interpolation interval, *N* is the number of nodes that the embedded structure consists of and 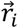 is the polar coordinate of the *i*th node. Value range of the vector angle is defined on (−π, π]. We used *l* = 100 to characterize *W*_*k*_ of CRC, T-ALL and corresponding normal controls with 40kb-resolution embedding results. For calculating *W*_*k*_ of embedded structures during early embryo developments of both human and mice, we used *l* = 10 for 40kb-resolution structures.

### Cluster displacement

For two sets of N-dimensional column vectors, to evaluate their difference in term of the ***i***th vector entry, the one-dimensional cluster displacement is defined as follows,

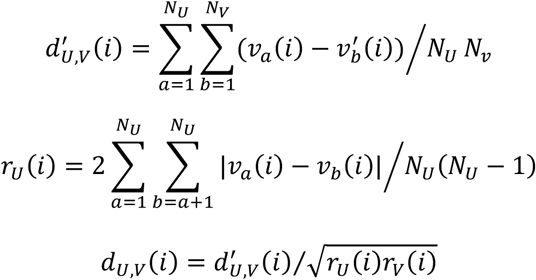

Here, *ν*(*i*) is the *i* th entry of vector *ν*, while *U* = {ν_1_, ν_2_, …, ν_a_} and 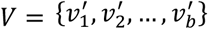 are two sets of vectors.

### Inter-chromosomal closeness matrix

We first calculated the average inter-chromosomal Poincaré distances for each pair of autosomes for each embedded structure,

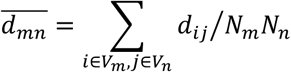

where *V*_*m*_ and *V*_*n*_ are sets of nodes belonging to chromosome *m* and *n* respectively, *d*_*ij*_ is the Poincaré distance between node *i* and *j, N*_*m*_ and *N*_*n*_ are the size of set *V*_*m*_ and *V*_*n*_ respectively. Each element in the inter-chromosomal distance matrix (22×22) were then divided by the geometric average of the corresponding pair of average intra-chromosomal distances, to evaluate the closeness between each chromosome pair.

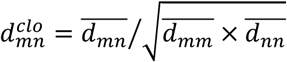

Each element in the closeness matrix was further normalized by subtraction of 1 and division by the maximum element, so that matrix values range from 0 to 1 and the intra-chromosomal closeness equals 0.

## Statistics

The significance levels of inequality of graph metrics between cancer/embryo samples and their corresponding normal controls were calculated under the one-sided Welch’s t-tests.

## Visualization methods

Statistical graphs and visualization of embedded CCNs in Poincaré disk were drawn by Matlab codes. Visualization of networks in schematics of Fig. 1 was performed by python package NetworkX.

## Data accessibility

The data that support the findings of this study were derived from the following resources available in the public domain: GEO and TCGA. The detailed data accession can be found in supplementary tables 1 and 2.

## Author Contributions

Conceptualization, YQ.G.; Data curation, JY.W., Y.X., YY.H., H.Q. and J.Z.; Formal analysis, JY.W., Y.X. and YY.H.; Funding acquisition, YQ.G.; Investigation, JY.W., Y.X., YY.H., H.Q. and J.Z.; Supervision, YQ.G.; Visualization, JY.W.; Writing – original draft, JY.W., Y.X., YY.H., H.Q. and J.Z.; Writing – review & editing, JY.W., Y.X. and YQ.G. All authors have read and agreed to the published version of the manuscript.

## Acknowledgments

The results shown here are part based upon data generated by the TCGA Research Network (http://cancergenome.nih.gov).

