## Supplementary information for "Characterization of network hierarchy reflects cell-state specificity in genome organization"

**Supplementary Figures 1-2**
**Supplementary Tables 1-4**

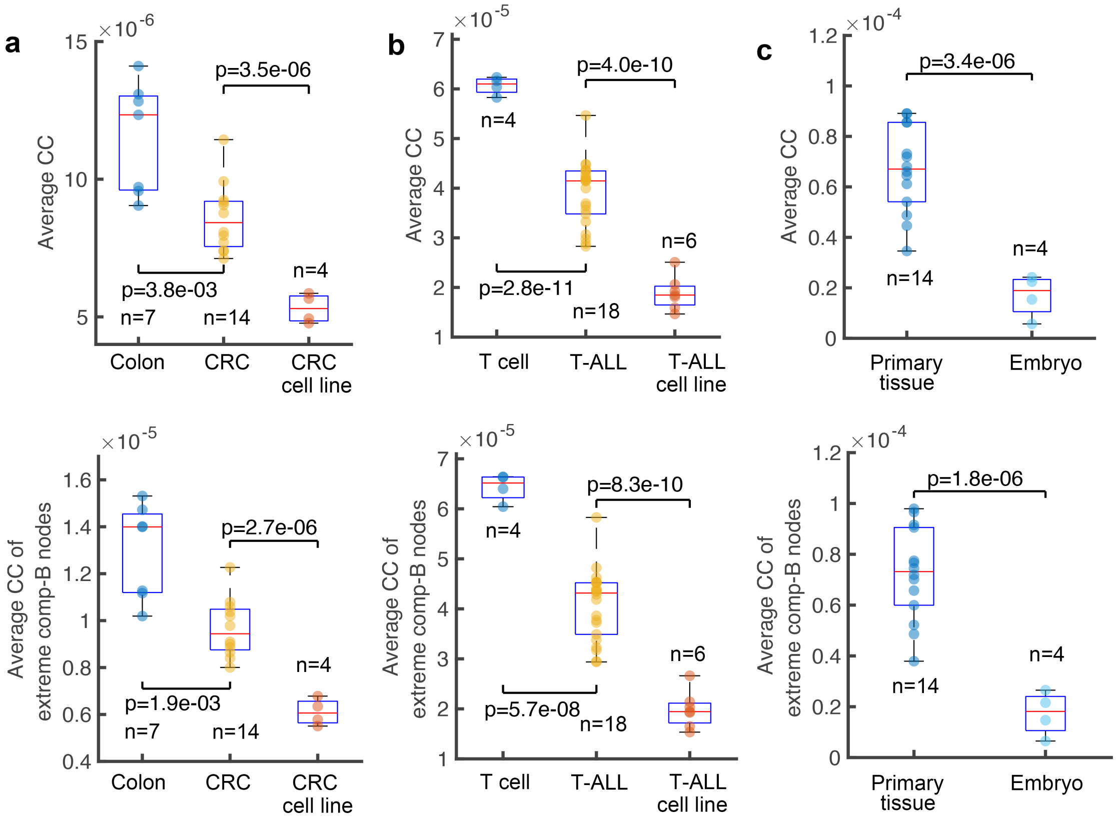

**Supplementary Fig. 1 Decrease of CC values in cancer and embryo CCNs compared to normal tissues.** Boxplots show average CC in cancer, cancer cell lines and corresponding normal samples for CRC (left) and T-ALL (middle). The right panel shows average CC of primary tissues and early embryos. For each sample, CC values were averaged among nodes in all intra-chromosomal CCNs. In the lower panel, CC values were averaged over nodes with the lowest 5% compartment vector entries on each chromosome.

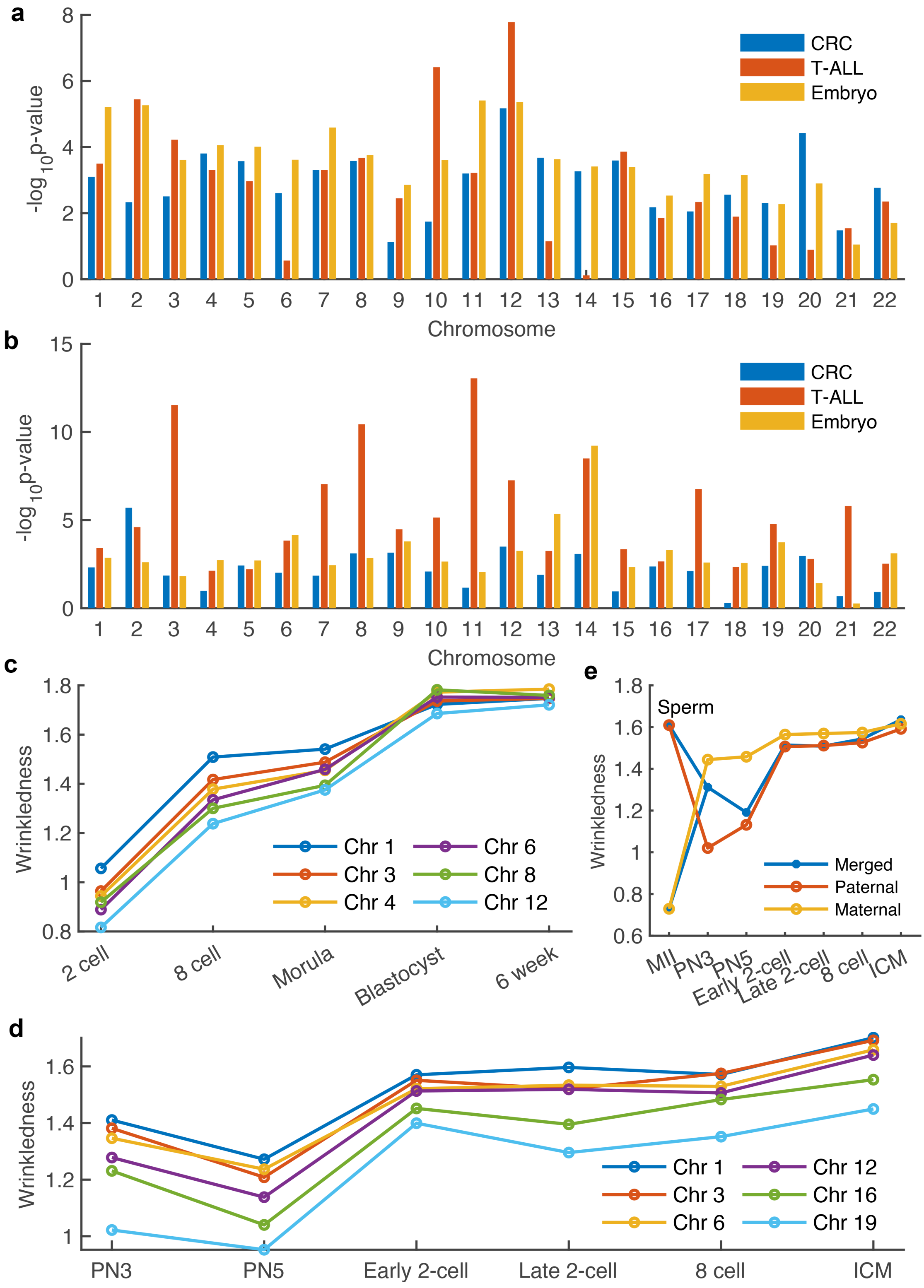

**Supplementary Fig. 2 Changes in hierarchy depths and structural wrinkledness in cancer and embryo development.** Vertical bars showing the significance of median $R_P$ increase (**a**) and wrinkledness ($W_k$) decrease (**b**) for each chromosome in three comparisons, which are CRC and T-ALL compared to their normal controls as well as embryos compared to primary tissues. Evolution of $W_k$ in early embryo developments of human (**c**) and mouse (**d**). **e** Average $W_k$ of all chromosomes for paternal and maternal alleles respectively as well as the merge of alleles in early embryo development of mouse.

**Supplementary table 1 Data information for Hi-C**

<table>
<tr><th>Cell type</th><th>Sample</th><th>Abbreviation</th><th>Description</th><th>Accession</th></tr>
<tr><td>Cortex</td><td colspan="2">CO</td><td rowspan="7">Normal tissue</td><td rowspan="7">GSE87112</td></tr>
<tr><td>Hippocampus</td><td colspan="2">HC</td></tr>
<tr><td>Left ventricle</td><td colspan="2">LV</td></tr>
<tr><td>Liver</td><td colspan="2">LI</td></tr>
<tr><td>Right ventricle</td><td colspan="2">RV</td></tr>
<tr><td>Small bowel</td><td colspan="2">SB</td></tr>
<tr><td>Spleen</td><td colspan="2">SX</td></tr>
<tr><td rowspan="7">Colon</td><td>BRD3328N</td><td>Colon 1</td><td rowspan="7">Adjacent normal tissues in resection specimens of CRC patients</td><td rowspan="24">GSE133928</td></tr>
<tr><td>BRD3409N</td><td>Colon 2</td></tr>
<tr><td>BRD3179N</td><td>Colon 3</td></tr>
<tr><td>BRD3462N</td><td>Colon 4</td></tr>
<tr><td>BRD3187N</td><td>Colon 5</td></tr>
<tr><td>BRD3170N</td><td>Colon 6</td></tr>
<tr><td>BRD3162N</td><td>Colon 7</td></tr>
<tr><td rowspan="12">Colon tumor</td><td>BRD3179</td><td>CRC 1</td><td rowspan="12">Colon tumors from CRC patients</td></tr>
<tr><td>MGH5328</td><td>CRC 2</td></tr>
<tr><td>BRD3412</td><td>CRC 3</td></tr>
<tr><td>MGH3535</td><td>CRC 4</td></tr>
<tr><td>BRD3462</td><td>CRC 5</td></tr>
<tr><td>BRD3187</td><td>CRC 6</td></tr>
<tr><td>BRD3162</td><td>CRC 7</td></tr>
<tr><td>BRD3378</td><td>CRC 8</td></tr>
<tr><td>MGH1904</td><td>CRC 9</td></tr>
<tr><td>MGH2231</td><td>CRC 10</td></tr>
<tr><td>MGH2834</td><td>CRC 11</td></tr>
<tr><td>MGH8416</td><td>CRC 12</td></tr>
<tr><td>Colon cell line</td><td colspan="2">FHC</td><td rowspan="5">Cell line</td></tr>
<tr><td rowspan="4">CRC cell line</td><td colspan="2">HCT116</td></tr>
<tr><td colspan="2">LS174T</td></tr>
<tr><td colspan="2">RKO</td></tr>
<tr><td colspan="2">SW480</td></tr>
<tr><td rowspan="4">Normal T cell</td><td>Normal 1</td><td>Normal T cell 1</td><td rowspan="4">Normal T-cells from peripheral blood</td><td rowspan="9">GSE146901</td></tr>
<tr><td>Normal 2</td><td>Normal T cell 2</td></tr>
<tr><td>Normal 3</td><td>Normal T cell 3</td></tr>
<tr><td>Normal 4</td><td>Normal T cell 4</td></tr>
<tr><td rowspan="5">Leukemia</td><td>076</td><td>T-ALL 1</td><td rowspan="5">T-lineage acute lymphoblastic leukemia (T-ALL)</td></tr>
<tr><td>077</td><td>T-ALL 2</td></tr>
<tr><td>093</td><td>T-ALL 3</td></tr>
<tr><td>097</td><td>T-ALL 4</td></tr>
<tr><td>098</td><td>T-ALL 5</td></tr>
</table>

| | | | | |
|---|---|---|---|---|
| | 102 | T-ALL 6 | | |
| | 103 | T-ALL 7 | | |
| | 107 | T-ALL 8 | | |
| | 108 | T-ALL 9 | | |
| | 115 | T-ALL 10 | | |
| | 116 | T-ALL 11 | | |
| | 117 | T-ALL 12 | | |
| | 118 | T-ALL 13 | | |
| | 121 | T-ALL 14 | | |
| | 122 | T-ALL 15 | | |
| | 123 | T-ALL 16 | | |
| | 124 | T-ALL 17 | | |
| | 032 | T-ALL 18 | | |
| T-ALL cell line | CEM | | Cell line | |
| | CUTLL1 | | | |
| | Jurkat | | | |
| | KE-37 | | | |
| | LOUCY | | | |
| | MOLT-16 | | | |
| | MOLT-4 | | | |
| 2-cell embryo | 2 cell | | Human early embryo | CRA000852 |
| 8-cell embryo | 8 cell | | | |
| Morula embryo | Morula | | | |
| Blastocyst embryo | Blastocyst | | | |
| 6-week embryo | 6 week | | | |
| Sperm | Sperm | | Generative cells of mouse | GSE82185 |
| MII | MII | | | |
| PN3 zygote | PN3 | | Mouse early embryo | |
| PN5 zygote | PN5 | | | |
| Early-2-cell embryo | Early 2 cell | | | |
| Late-2-cell embryo | Late 2 cell | | | |
| 8-cell embryo | 8 cell | | | |
| ICM | ICM | | | |

**Supplementary table 2 Data information for RNA-seq of CRC**

| Sample | Description | Accession |
|---|---|---|
| TCGA-A6-2671 | The sample ID followed by "-11A" or "-01A" is the normal tissue adjacent to tumor or the tumor tissue of the patient. | TCGA |
| TCGA-A6-2675 | | |
| TCGA-A6-2678 | | |
| TCGA-A6-2679 | | |
| TCGA-A6-2680 | | |
| TCGA-A6-2682 | | |

| | | |
|---|---|---|
| TCGA-A6-2683 | | |
| TCGA-A6-2684 | | |
| TCGA-A6-2685 | | |
| TCGA-A6-2686 | | |
| TCGA-A6-5659 | | |
| TCGA-A6-5662 | | |
| TCGA-A6-5665 | | |
| TCGA-A6-5667 | | |
| TCGA-AA-3489 | | |
| TCGA-AA-3496 | | |
| TCGA-AA-3511 | | |
| TCGA-AA-3514 | | |
| TCGA-AA-3516 | | |
| TCGA-AA-3517 | | |
| TCGA-AA-3518 | | |
| TCGA-AA-3520 | | |
| TCGA-AA-3522 | | |
| TCGA-AA-3525 | | |
| TCGA-AA-3527 | | |
| TCGA-AA-3531 | | |
| TCGA-AA-3534 | | |
| TCGA-AA-3655 | | |
| TCGA-AA-3660 | | |
| TCGA-AA-3662 | | |
| TCGA-AA-3663 | | |
| TCGA-AA-3697 | | |
| TCGA-AA-3712 | | |
| TCGA-AA-3713 | | |
| TCGA-AZ-6598 | | |
| TCGA-AZ-6599 | | |
| TCGA-AZ-6600 | | |
| TCGA-AZ-6601 | | |
| TCGA-AZ-6603 | | |
| TCGA-AZ-6605 | | |
| TCGA-F4-6704 | | |
| BRD3187N | Normal tissue adjacent to tumor | GSE133928 |
| BRD3162 | Tumor | |
| BRD3174 | | |
| BRD3179 | | |
| BRD3187 | | |
| MGH2834 | | |
| MGH5328 | | |

| MGH8416 |
|---|

**Supplementary table 3 Gene set enrichment analysis of genes in intersected bridge ends in normal colons and CRCs**

| Gene Ontology ID | Description | p-value | FDR |
|---|---|---|---|
| GO:0002673 | regulation of acute inflammatory response | 1.40E-05 | 0.04566856 |
| GO:0006958 | complement activation, classical pathway | 2.95E-05 | 0.04566856 |
| GO:0030449 | regulation of complement activation | 3.00E-05 | 0.04566856 |
| GO:2000257 | regulation of protein activation cascade | 3.83E-05 | 0.04566856 |
| GO:0002455 | humoral immune response mediated by circulating immunoglobulin | 8.40E-05 | 0.08021057 |
| GO:0002526 | acute inflammatory response | 0.00011602 | 0.09231038 |
| GO:0006956 | complement activation | 0.00030326 | 0.20682222 |
| GO:0070613 | regulation of protein processing | 0.0004441 | 0.26501863 |
| GO:0002920 | regulation of humoral immune response | 0.0005226 | 0.27711422 |
| GO:1903317 | regulation of protein maturation | 0.00058047 | 0.27711422 |
| GO:0002921 | negative regulation of humoral immune response | 0.00173265 | 0.75197111 |

**Supplementary table 4 Gene set enrichment analysis of genes with Z-score of $R_p$ smaller than -10 in CRCs compared to small bowel**

| Gene Ontology ID | Description | p-value | FDR |
|---|---|---|---|
| GO:0090023 | positive regulation of neutrophil chemotaxis | 7.30E-07 | 0.00344539 |
| GO:0071624 | positive regulation of granulocyte chemotaxis | 2.69E-06 | 0.00423098 |
| GO:1902624 | positive regulation of neutrophil migration | 2.69E-06 | 0.00423098 |
| GO:0090022 | regulation of neutrophil chemotaxis | 5.80E-06 | 0.00684355 |
| GO:1902622 | regulation of neutrophil migration | 1.63E-05 | 0.01534759 |
| GO:0071622 | regulation of granulocyte chemotaxis | 0.00036237 | 0.28500329 |
| GO:0048278 | vesicle docking | 0.0011462 | 0.77270277 |
| GO:0070493 | thrombin-activated receptor signaling pathway | 0.00174944 | 0.9770344 |
| GO:0006904 | vesicle docking involved in exocytosis | 0.00221446 | 0.9770344 |
| GO:0031640 | killing of cells of other organism | 0.00240671 | 0.9770344 |
| GO:0044364 | disruption of cells of other organism | 0.00240671 | 0.9770344 |
| GO:0060192 | negative regulation of lipase activity | 0.00265313 | 0.9770344 |
| GO:0002690 | positive regulation of leukocyte chemotaxis | 0.00269156 | 0.9770344 |
